# Predictive feedback to V1 dynamically updates with sensory input

**DOI:** 10.1101/180539

**Authors:** Grace Edwards, Petra Vetter, Fiona McGruer, Lucy S. Petro, Lars Muckli

## Abstract

Predictive coding theories propose that the brain creates internal models of the environment to predict upcoming sensory input. Hierarchical predictive coding models of vision postulate that higher visual areas generate predictions of sensory inputs and feed them back to early visual cortex. In V1, sensory inputs that do not match the predictions lead to amplified brain activation, but does this amplification process dynamically update to new retinotopic locations with eye-movements? We investigated the effect of eye-movements in predictive feedback using functional brain imaging and eye-tracking whilst presenting an apparent motion illusion. Apparent motion induces an internal model of motion, during which sensory predictions of the illusory motion feed back to V1. We observed attenuated BOLD responses to predicted stimuli at the new post-saccadic location in V1. Therefore, pre-saccadic predictions update their retinotopic location in time for post-saccadic input, validating dynamic predictive coding theories in V1.

## Introduction

Predictive coding accounts of vision propose that higher cortical areas use internal models of the world to predict sensory inputs (Bastos et al., 2012; Clark, 2013; Friston, 2005; Mumford, 1992; Rao and Ballard, 1999). Cortical feedback carries these predictions back to V1 where a neural mechanism compares them to the actual sensory inputs (see Alink et al., 2010). However, a central tenet of predictive coding remains to be tested. Humans saccade approximately three times per second, changing the retinotopic pattern of sensory inputs to V1 (Adams et al., 2007). Therefore, for cortical predictive feedback to be functional, it must update to new retinotopic locations in V1 in time to meet post-saccadic input (Melcher, 2011; Mumford, 1991).

Central to our study is the creation of an internal model in the brain, during which feedback carries sensory predictions from higher areas down to V1. The apparent motion illusion offers a paradigm for investigating such a model. Apparent motion is an illusion of a moving token between two alternating flashing stimuli (KOLERS, 1963; Shepard and Zare, 1983). Apparent motion is integrated in V5 (Akselrod et al., 2014; Muckli et al., 2002; Sterzer et al., 2006a; Vetter et al., 2015; Wibral et al., 2009), which feeds back a retinotopically-specific spatiotemporal prediction of a moving token to V1, inducing activation along the non-stimulated illusory motion trace (Ahmed et al., 2008; Alink et al., 2010; Larsen et al., 2006; Muckli et al., 2005; Sterzer et al., 2006a). The spatio-temporal prediction on the apparent motion trace can be probed with targets presented either in-time with the motion illusion (predicted), or out-of-time (unpredicted). In-time targets are detected more accurately than out-of-time targets (Schwiedrzik et al., 2007; Vetter et al., 2015, 2014, 2012) but lead to less (i.e. dis-amplified) BOLD activation in V1 compared with amplified activation for out-of-time targets(Alink et al., 2010). According to predictive coding theories, the amplified BOLD activation for out-of-time targets is indicative of an error signal because they are less predictable in the context of the illusory moving token. Likewise, dis-amplified BOLD activity for in-time targets is consistent with the successful prediction (or ‘explaining away’) of sensory input. Here, we tested if pre-saccadic predictions feed back to new retinotopic locations in V1 in time for post-saccadic target processing. To this end, we presented the apparent motion illusion to one visual hemifield and then prompted a saccade, transferring motion-related feedback to the opposite hemisphere. Our data confirms that cortical feedback dynamically updates to new predicted retinotopic coordinates in V1.

## Results

We first induced an apparent motion illusion, which creates a spatio-temporally precise prediction of apparent motion along the illusory motion trace in V1. We then instructed subjects to perform a right-to-left saccade across the apparent motion stimulus. When the saccade landed left of the apparent motion illusion, a target was presented either in-time or out-of-time during the last cycle of apparent motion (Figure 1a-d). After the saccade, the stimulus was in the right visual field, and processed by post-saccadic left V1. The post-saccadic left V1 processed only one apparent motion inducer after the target appeared, thus illusion-related activity is present even though the post-saccadic left V1 was not stimulated with a full apparent motion cycle. Without prior expectation from the contralateral hemisphere, both target stimuli are equally predictable by the surrounding stimulation. We tested for the presence of post-saccadic predictions by comparing the BOLD activity to in-time and out-of-time targets, in a test region on the illusory motion trace in post-saccadic left V1 (Figure 2). This test region corresponds to the new retinotopic location at which the cortex processes the target stimulus after the saccade. We examined this test region in three apparent motion conditions: no target, in-time target, and out-of-time target. Subjects were instructed to report if they perceived a target presented immediately after saccade. We ran four functional magnetic resonance imaging (fMRI) experiments. Our pilot fMRI experiment led us to use high-resolution eye-tracking data for trial exclusion in the main apparent motion experiment. In the pilot experiment, we found consistent results with the main experiment in the subjects with valid eye movement data (see supplemental material and Figure S1). To confirm that our results are due to predictions created by the internal model of apparent motion, we also ran a flicker control fMRI experiment. Here, we presented the two flashing inducer stimuli simultaneously, instead of in alternating rhythm as in the apparent motion stimulation (Figure 1e). In the context of the flicker stimulation, neither target had a temporal predictability but for simplicity we kept the conditions labeled as “in-time” and “out-of-time” as the targets were presented at the same stimulus onset time in relation to the lower apparent motion inducer stimulus as in the main fMRI experiment. Finally, we ran a replication of our main apparent motion and flicker experiments. The design of the replication was identical to the main and flicker experiment except for two variations: (1) we ran the replication as a within group design and (2) we used the same hardware throughout the replication (32-channel head coil) while in the original experiment we switched between experiments using the 12-channel head coil for the main and the 32-channel for the flicker control. The replication supported and strengthened our original findings.

**Figure 1:**
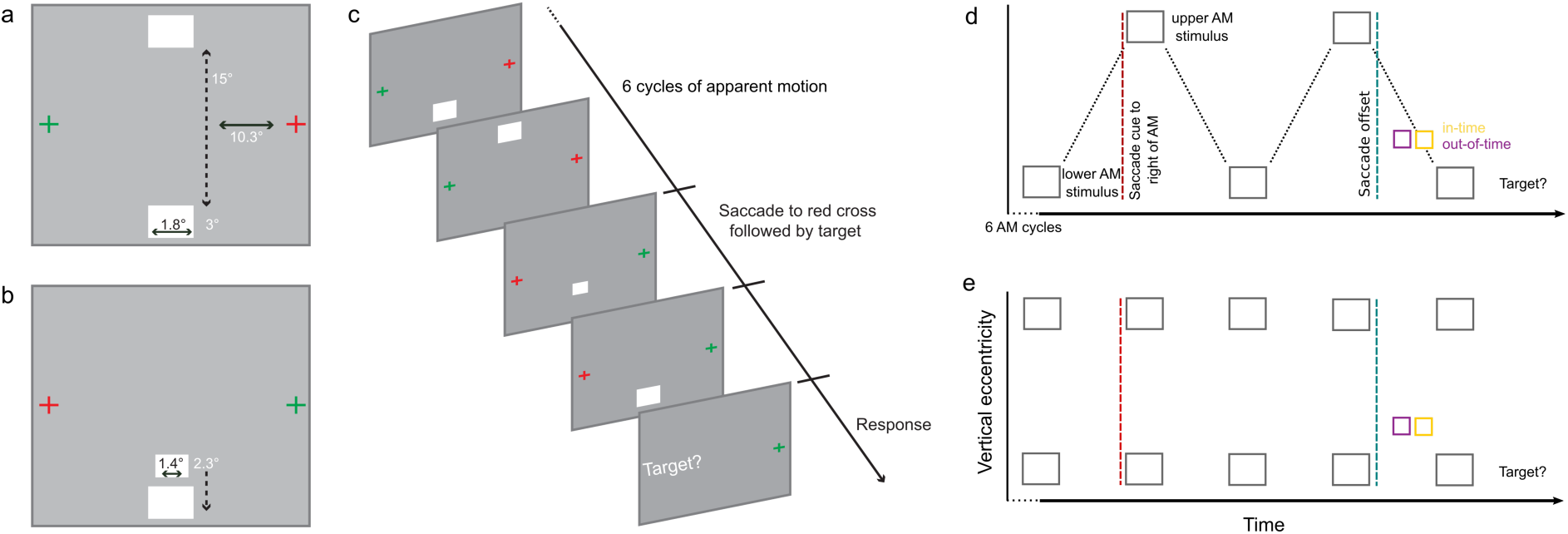
Apparent Motion Stimuli. **(a)** Stimulation before saccade: Subjects fixated on the red cross. Apparent motion inducing stimuli flashed in alternation to the left of fixation. **(b)** Stimulation after saccade: subjects fixated on the red cross, with the motion trace now presented to the right of fixation and processed by the left hemisphere. Immediately after saccade landing on the red cross, we presented a target on the apparent motion trace either in-time or out-of-time with the illusion. Visual angles are shown for main experiment (rounded). **c)** Diagrammatic apparent motion trial. **d)** Apparent motion stimulus timeline. The red cross was located to the right of the apparent motion stimulus for 6 cycles. Within each cycle, we presented the upper apparent motion inducer for 83.3ms, followed by an inter-stimulus interval of 83.3ms, and then the lower apparent motion inducer for 83.3ms and another inter-stimulus interval of 83.3ms before the next cycle began. The red fixation cross moved to the left of the apparent motion, cuing saccade prior to cycle seven, depicted by the red dashed line. During valid trials, the saccade landed left of the apparent motion during the 8^th^ cycle (blue dashed line), and a target was presented either in-time or out-of-time. We presented targets for the duration of one frame (16.67ms) 450ms or 483.3ms after the saccade cue. We only accepted trials in which subjects executed the saccade within 400ms after the cue (but not before 333ms after the cue). Therefore we presented in-time targets at least 83.3ms after saccade offset, and out-of-time targets at least 50ms after saccade offset. Note that the post-saccadic stimulated left hemisphere saw only one inducer after the target appeared. After the illusion ceased, we prompted subjects to respond if they detected a target or not. **e)** Flicker trial timeline. The flicker trials are the same as the apparent motion trials, but with upper and lower inducers presented simultaneously.

### Post-saccadic apparent motion prediction at the new retinotopic location in left V1

We tested for prediction-related activity on the illusory motion trace by first mapping the cortical representation of the target stimulus located between the two blinking apparent motion inducing stimuli. We presented a static target stimulus in between the apparent motion inducers, and a static stimulus at the same location as the lower apparent motion inducer. To map the target region of interest (ROI), we used a GLM contrast of ‘target stimulus’ > ‘lower AM inducer stimulus’ (Figure 2, online methods). After identifying the target ROI in post-saccadic left V1, we compared activation for the three apparent motion conditions. The illusory trace was activated during apparent motion even when no target was presented (no-target condition) in 5/9 subjects for the Main experiment, and in 7/8 subjects for the Replication experiment (single subject: *p<0.05; Main experiment group mean beta-weight (SE)=0.2(0.02), p<0.0001, Replication group mean beta-weight (SE) = 1.9(0.3), p=0.0007 Figure 3a-b & Figure 4a & 4b). Importantly, we observed apparent-motion trace activity in post-saccadic (left) V1 even though we only presented one apparent motion inducing stimuli to this hemisphere. Illusion-related activity on the non-stimulated apparent motion trace has been shown before (Ahmed et al., 2008; Larsen et al., 2006; Muckli et al., 2002, 2005; Sterzer et al., 2006a), but it is novel to find that the illusory activity transfers across hemispheres to the post-saccadic (left) V1 in anticipation of post-saccadic continuation of the illusion. It is unlikely that activity found for the ‘no target apparent motion’ condition is due to the lower apparent motion stimulus. We conservatively selected voxels specific to the target-processing region and confirmed their retinotopic position on individual maps. At the target ROI, activity to apparent motion was higher than to the mapping stimulus confirmed by a GLM contrast between ‘no target apparent motion’ and the ‘lower inducer’ (Main experiment: t(8)=3.81,p=0.005, Replication: t(7)=2.98,p=0.02). Therefore, at the target position it is the illusion which drives activity, not the inducer stimulus. In contrast, at the inducer position, activation for the mapping stimulus was stronger than the apparent motion stimulus, which we expect because the mapping stimulus was shown 48 times longer than the apparent motion inducing stimulus (4000 ms vs 83.3 ms). Lastly, individual subject activity maps indicate that the peak apparent motion activation overlaps more with the target region than with the inducer stimulus (Figure 4a & 4b).

**Figure 2:**
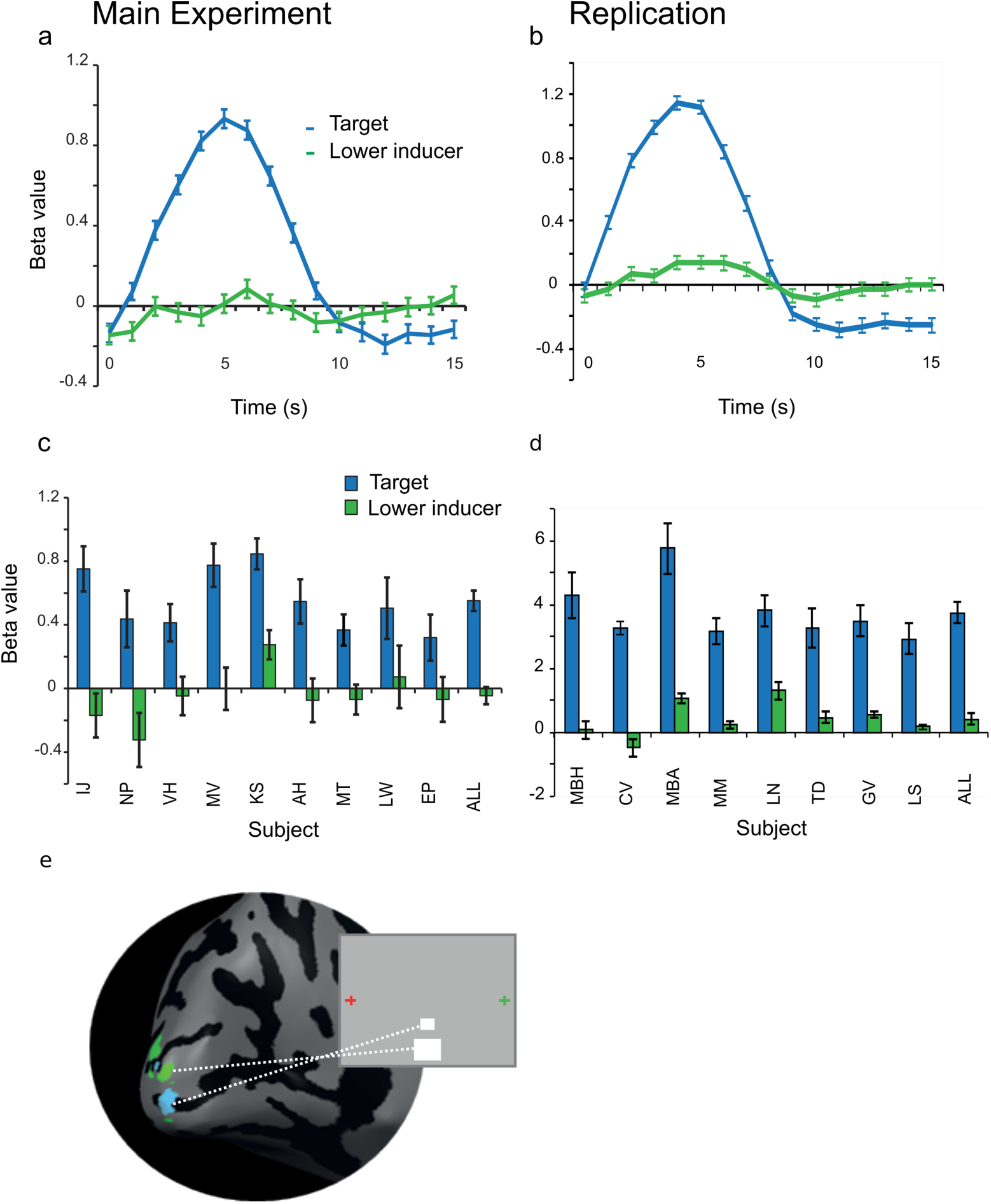
Retinotopically Defined Target Region of Interest. The target ROI (blue) responds specifically to the target and not to either AM inducer. Left hand column corresponds to data from main experiment, and right column to data from the replication of the main apparent motion experiment. **(a) & (b)** Group event-related BOLD responses to the target and lower inducer stimuli, in the target region of interest (mean ± SEM). **(c) & (d)** Single subject and group beta-weight values averaged over the peak BOLD activation to the target and lower inducer, in the target region of interest, in the main experiment (mean ± SEM; *p<0.05). **(e)** Cortical representation of the target stimulus (blue) and the lower inducer stimulus (green), on the inflated left cortical hemisphere of one subject.

To investigate apparent motion prediction in post-saccadic (left) V1, we tested whether in-time and out-of-time targets were dis-amplified or amplified (Figure 3b & 3c). If predictive codes update to new retinotopic positions in post-saccadic (left) V1, in-time target activation may be dis-amplified (because targets are more predictable) and out-of-time targets amplified(Alink et al., 2010) (because targets are less predictable). We found less increased BOLD in the left V1 target ROI for in-time versus out-of-time targets in 7/9 subjects for the Main experiment (group mean (SE) beta-weight for in-time target trials = 0.23(0.13); out-of-time target trials = 0.29(0.17); t(8)=2.39, p=0.04, Figure 3c) and in 5/8 for the Replication (group mean (SE) beta-weight for in-time target = 1.99(0.35); out-of-time target trials = 2.3(0.33, t(7)2.62,p=0.035, Figure 3d)). The increase of BOLD activity for out-of-time targets may indicate a prediction error signal or dis-amplification for predicted in-time targets. This error signal occurs at the new retinotopic location due to pre-saccadic predictions updating their retinotopic location in time for post-saccadic input.

**Figure 3:**
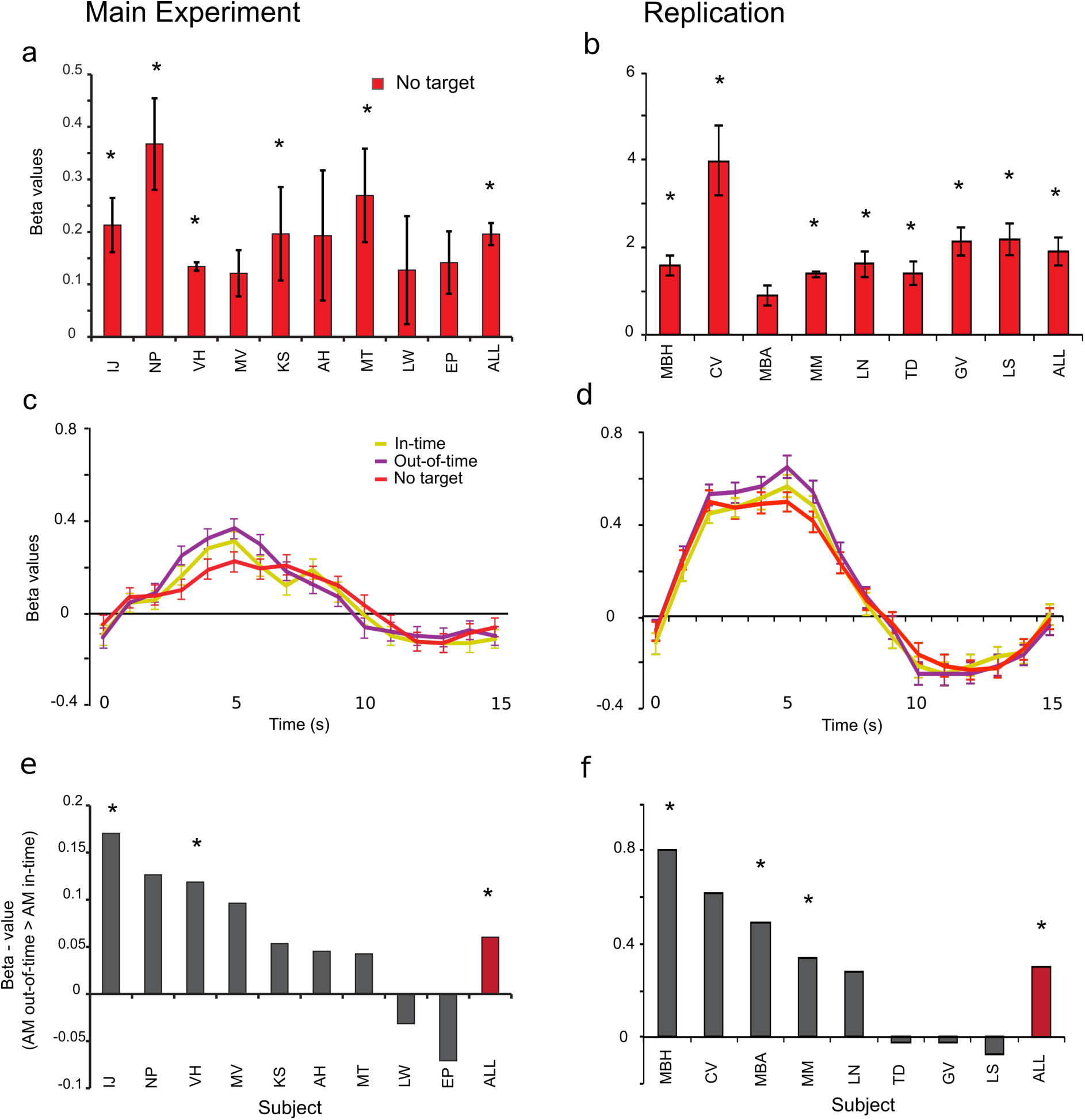
BOLD Response to Post-Saccadic Apparent Motion for Main fMRI experiment (left) and Replication (right) in Left Hemisphere V1. **(a) & (b)** Difference in beta-weight values of ‘apparent motion with no target’ greater than ‘baseline’ conditions, in the target region. Beta-weight values were averaged over the peak activation (*p<0.05) for single subjects and the group (mean ± SEM; main experiment: n=9, replication: n=8). **(c) & (d)** Group-averaged event-related BOLD responses to all apparent motion conditions in the target region (mean ± SEM). **(e) & (f)** Difference between in-time and out-of-time target trials for single subjects and the group (red bars) in the main apparent motion experiment & replication. The control and replication experiments used a 32-channel coil which provides higher functional activation magnitudes in visual cortex than the 12-channel coil used in the main experiment (Kaza et al., 2011).

With the flicker control experiment we show that the increased BOLD activity for out-of-time targets is specific to the spatiotemporal expectation triggered by the apparent motion context. For the flicker control experiment, we again mapped the retinotopic region of left V1 processing the target stimulus but we found, as expected, no significant difference between the in-time and out-of-time targets in the group analysis (non-parametric signed rank test: Main experiment: p=0.625, figure 5a; Replication: t(7)0.35, p=0.73, figure 5b). Only one subject showed a significant difference between in-time and out-of-time target activity in the Replication flicker experiment, with an increased activity for in-time targets (p<0.05). We tested if the difference in activation to in-time and out-of-time targets was greater in the main apparent motion experiment (where we expect an enhanced response to the out-of-time target) than in the flicker control (where we expect to find no consistent pattern of difference in activation to the target conditions). We conducted a t-test which does not assume equal variance or equal group size for the main fMRI experiment versus the flicker experiment and found no significant activity difference between ‘in-time’ and ‘out-of-time’ targets (p=0.3699). When we replicated our apparent motion experiment and flicker control using the within subjects design, we found that activation differences between in-time and out-of-time targets were significantly different between the apparent motion stimulation and the flicker stimulatio (t(7)2.65, p=0.03). Our use of a within subject design for the replication study reduced the variability and increased power of analysis. When pooling across all experiments, a rank sums test on unequal groups between AM (n=22) and flicker (n=13) reveals that activation differences between in-time and out-of-time targets are significantly different (p=0.04). Flicker perception might trigger different low precision predictions for additional target appearance in different individuals explaining the higher variance we see in subjects’ BOLD responses. In summary, the pattern of results show, that only the internal model of apparent motion imposes a spatiotemporal expectation on the processing of the target. Without the internal model of apparent motion illusion, one temporal sequence (i.e. target stimulus) cannot be more expected than the other, and we observe no activity difference in target processing. This also confirms the timing of the targets relative to the inducer does not create a differential BOLD signal by itself, but only if it is accompanied by the internal model of an apparent motion illusion.

### Location and temporal specificity of apparent motion prediction in V1

We find the internal model of apparent motion (but not flicker) creates a predictive signal at the post-saccadic target processing location in left V1. To ensure this signal is spatially and temporally specific, we ran a 3-way ANOVA on location in V1 (target ROI and lower inducer ROI) vs. time-point of trial (pre-saccade and post-saccade) vs. condition (in-time or out-of-time target activity). We found a significant interaction between location, target condition and time-point for our Main experiment (F(1,8)6.46,p=0.035; Figure 5c) and for our Replication (F(1,7)7.9,p=0.007; Figure 5d). Post-hoc tests revealed the interaction is driven by time-point (p<0.04) and ROI (p<0.05), the out-of-time versus in-time difference only reported at the post-saccadic time-point accounts for these findings in both the Main (p=0.04) and Replication (p=0.035) experiments. Note, there was no significant difference between in-time and out-of-time targets at the post-saccadic time-point in the lower inducer stimulus processing region for the Main (p>0.48) or Replication experiments (p>0.14). Prior to the saccade we found no significant difference between conditions in neither the target nor the apparent motion region (p>0.05). This supports our interpretation that trans-saccadic cortical feedback to V1 is spatially and temporally specific to the region of cortex processing a predicted sensory input. Furthermore, in figure 4c and 4d we present individual subject cortical inflations demonstrating that voxels activated for the predictive effect (out-of-time > in-time target activity) overlap and surround the cortical representation of the target stimulus but rarely spill over into region of cortex processing the lower inducer stimulus.

**Figure 4:**
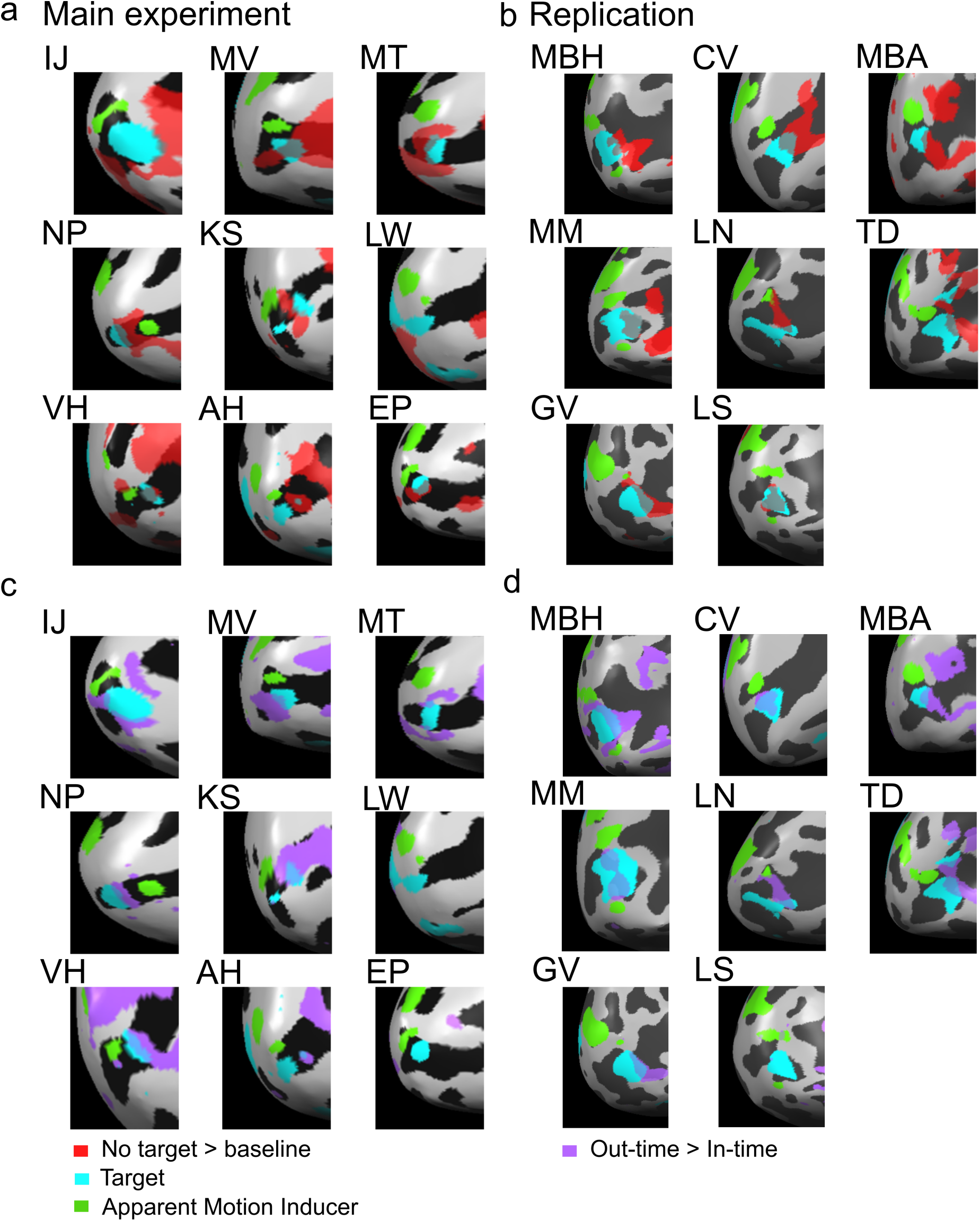
Individual Subjects’ BOLD Activation Along the Calcarine Sulcus in Left Hemisphere V1. Main fMRI experiment subjects left, replication subjects right. **(a) & (b)** No target Apparent motion > baseline (red) shown on the inflated left occipital cortex for all nine subjects. Each map also depicts the retinotopically-mapped regions of interest (ROIs). The target ROI is presented in blue and the lower apparent motion inducer ROI in green. No target apparent motion activity is located on and around the target ROI, rarely spilling onto the lower apparent motion inducer ROI. **(c) & (d)** Out-of time > in-time target activity (purple) shown on inflated left occipital cortex for each subject. As with (4a), out-of-time > in-time target activity is found within and around the target ROI.

**Figure 5:**
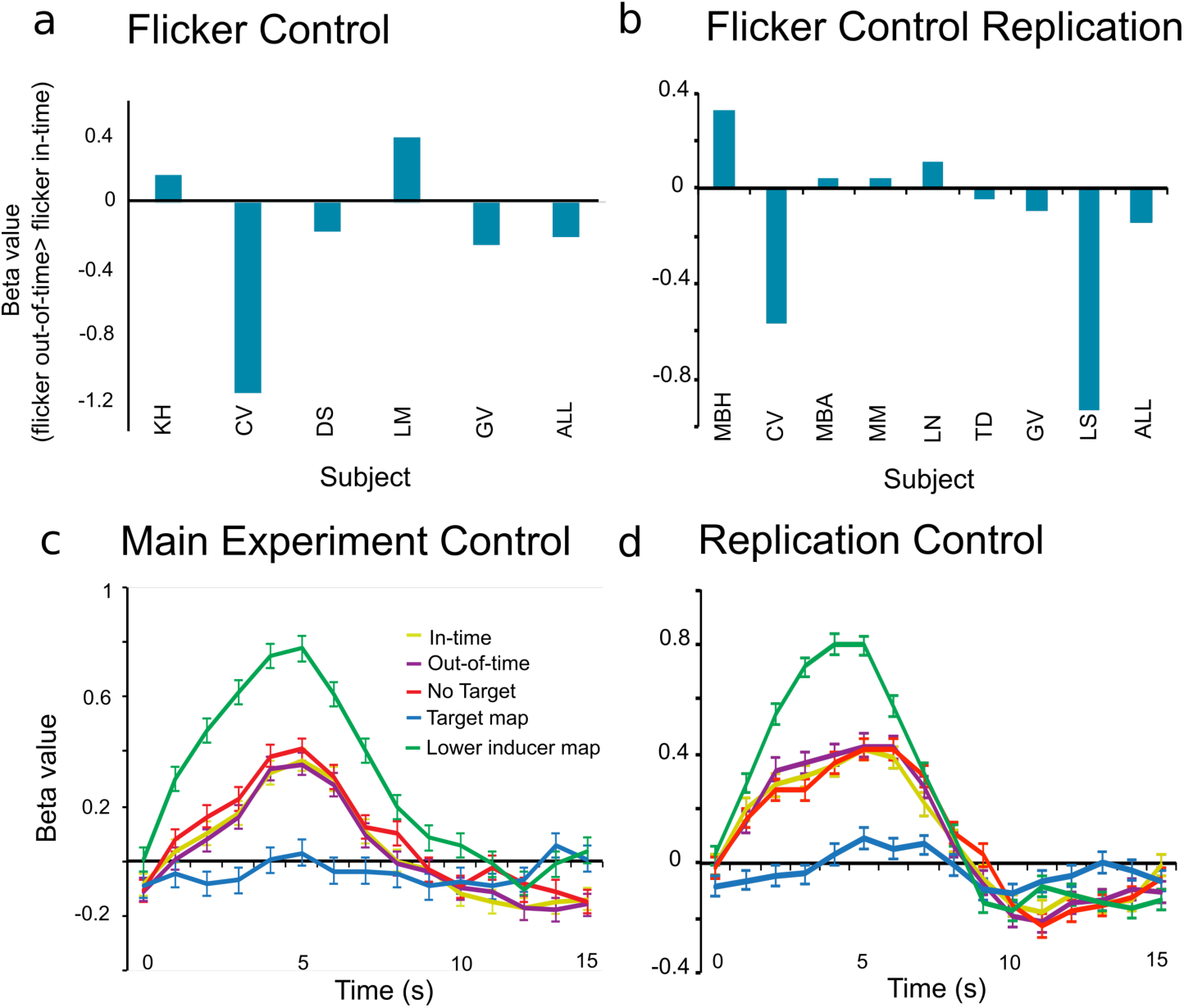
Control Data. Main experiment data in left column, replication in right column. **(a) & (b)** BOLD Response to Flicker Control: No significant difference in beta-weight values between in-time and out-of-time target trials for single subjects and the group in the target region in the flicker control experiments. **(c) & (d):** Group-averaged event-related BOLD responses to all apparent motion conditions at the apparent motion inducer region (contrast: Lower inducer (green) > target (blue)).

### Region specific apparent motion prediction

In the region of left dorsal V2 activated by the target stimulus, we found weak responses to all apparent motion conditions after the saccade for both Main and Replication experiments (in-time targets p<0.05; out-of-time targets p<0.05; no target p<0.003; Figure S2a & S2b). We also confirmed activation of motion-sensitive right and left V5 during apparent motion perception (Figure S2c & S2d). Right and left V5 were mapped using a contrast of apparent motion with no target > baseline. We find a significant effect of region when performing a repeated measures ANOVA for the out-of-time>in-time activation difference across all regions (left V1, left V2, right V5 and left V5) in the main experiment (F(3,8) 2.97, p=0.03) and replication (F(3,7) 3.77, p<0.01). We do not find the out-of-time > in-time target effect in V2 (Main: p=0.57; Replication: p=0.87), right V5 (Main: p=0.3; Replication: p=0.45), or left V5 (Main: p=0.9; Replication: p=0.85) in either experiment, meaning that our ANOVA is driven by the out-of-time>in-time target effect in V1 reported previously (Main: p=0.04; Replication: p=0.035). Our lack of prediction effect in left and right V5 is consistent with previous findings (Alink et al., 2010).

Since V2 has larger receptive fields, the target ROI also responded slightly to the lower AM inducing stimulus (Main experiment: beta-weight=0.16(0.05), p<0.009; Replication: beta-weight=0.41(0.046), p<0.001; Figure S2a & S2b). The lack of retinotopic specificity might have affected the sensitivity to pick up differential effects amongst the AM conditions. Alternatively, it might indicate a stronger prediction effect in V1 than in V2.

### Extra-session psychophysical control experiment

During the fMRI experiments, we found no behavioral detection advantage for either in-time or out-of-time targets as shown previously (Figure S4; (Schwiedrzik et al., 2007; Vetter et al., 2012). In a separate behavioral experiment, subjects were again required to respond if they detected a target presented within the apparent motion trace after saccade (Online Methods). Subjects were more accurate at detecting predictable in-time targets after saccades. The in-time advantage effect was significant at group level (t(7)3.07,p=0.015) and for 4/8 subjects (Figure S4c; p<0.02), replicating previous work with fixation (Schwiedrzik et al., 2007; Vetter et al., 2015) and after eye-movement (Vetter et al., 2012).

## Discussion

We provide evidence for dynamic updating of cortical predictions to new retinotopic locations at the lowest level of the cortical hierarchy: primary visual area V1. Our findings support the proposals of Helmholtz (see Cahan, 1993) and Gregory (1966) that the brain creates predictive models explaining away expected changes, including those made by planned eye-movements. Moreover our results support the theories of Mumford (1992) and Friston (2010) who suggest visual predictions descend the cortical hierarchy in feedback. However, until now, it has not been established that cortical predictions feed back to retinotopic visual cortex in time to interact with post-saccadic sensory input. We measured fMRI of subjects presented with the apparent motion illusion, during which V5 feeds back predictions to V1. We then prompted subjects to make a saccade, necessitating that predictive feedback from V5 updates to a new retinotopic location in V1. In our data, we observed a top-down signal in V1 which is retinotopically-precise, predictive of spatiotemporal sensory inputs and effective after a saccade. We confirm an anticipated, but as yet unobserved, mechanism of human vision: cortical predictions update to new locations in retinotopic cortex in time for post-saccadic processing.

### Prediction in vision

The world is not deconstructed each time we move our eyes, and reconstructed after each new fixation. Instead, the brain maintains perceptual continuity (e.g. Fischer and Whitney, 2014), because visual experience is the result of the brain’s internal model of the physical world merged with sensory signals (Lee et al., 1998). The brain performs this modeling process in order to operate most energy-efficiently, and eye movements function to test these internal models (Friston et al., 2012). Under predictive coding hypotheses, internal models generate predictions of forthcoming sensory inputs (Fairhall et al., 2017). Cortical feedback carries these predictions down the visual hierarchy where a neural mechanism compares them with actual sensory inputs. However, the early visual system is flooded with different patterns of sensory inputs each time we move our eyes(Adams et al., 2007). This means that if the cortex is to compare predictions to sensory inputs, predictions must update to the new retinotopic location in sensory cortex. This study is the first to provide evidence for the retinotopic updating of predictive feedback to V1.

### Predictive cortical feedback updates to new retinotopic locations with long-range saccades

In V1, cortical predictions must be spatially and temporally specific in order to be functional. Our spatial location of interest was a region of V1 that responds to a portion of the visual field between two apparent motion inducing stimuli. It was in this region of cortex that we observed a lower BOLD response to predictable versus unpredictable targets. This attenuation in BOLD signal is due to V5 feeding back predictions (Vetter et al., 2015) which dis-amplify expected inputs in sensory cortex (Alink et al., 2010). Over the trial time there are cognitive and feedforward effects that may contribute to the observed activation in the post-saccadic apparent motion trace. Our subjects observed apparent motion initially in the left visual hemifield. During this time we assume attention is divided between the red fixation cross in the fovea, the illusion percept (presented peripherally), and the future fixation position (green fixation cross, even further peripherally). During this pre-saccadic phase, we hypothesize three ongoing processes: (1) anticipatory remapping of the apparent motion stimulus due to the upcoming saccade (2) the establishment of visual predictions in higher areas induced by apparent motion, and (3) task related, top-down attention processes monitoring color changes in the peripheral location. In the post-saccadic phase, we observed in left V1: apparent motion related activity, on top of which we found target-related activity that is amplified if out-of-time with the apparent motion illusion (or dis-amplified in case of a spatiotemporal match between the prediction of illusory motion and the in-time target).

Let us turn first to how saccade-related remapping signals could modulate our target region in V1. Receptive field remapping is the anticipatory activation of receptive fields which are about to receive sensory input after saccade (Melcher and Colby, 2008). In our ‘no-target’ condition, the activity that we observed on the apparent motion trace in the left V1, i.e. at the new retinotopic location could be due to remapping effects in this region of cortex (see Merriam et al., 2007). Apparent motion in the left hemifield induces an activity increase on the apparent motion trace in the right hemisphere (Muckli et al., 2005) and we suggest this activity likely remaps to the new, left hemisphere. However, for both the in-time and out-of-time target activity on the apparent motion trace, receptive field remapping is not a credible explanation because we did not present a target in the left hemifield to be remapped. Further, although remapping activity might be present around the AM inducers and the fixation cross, these stimuli all map to regions of cortex separate from that processing the AM trace. Lastly, if anticipatory activity related to upcoming saccade selection accounted for the activation we observed on the apparent motion trace during the no-target conditions, it would not explain the difference between in-time and out-of-time targets.

The next top-down influence we consider is spatial attention. How does visual spatial attention contribute to our observed activation pattern on the post-saccadic apparent motion trace? A recent study by Cravo et al., demonstrated increased target detection during low-frequency phase entrainment using rhythmic stimuli. The authors suggest oscillations carry temporal expectations during attention (Cravo et al., 2013). However, our main effect is the difference in BOLD activation for in-time and out-of-time targets, which attention models struggle to account for in two ways. Attention is often conceptualized as a gain model where increased activity correlates with better performance (Brefczynski and DeYoe, 1999; Gandhi et al., 1999). In our findings, out-of-time targets induce increased BOLD activity but in an extra-session psychophysical we replicated earlier findings consistently showing subjects detect them less accurately than in-time targets. Moreover, even putting these limitations aside, we have shown previously that when subjects direct attention away from the location of apparent motion, feedback still fills in a “moving” token on the apparent motion trace (Muckli et al., 2005).

We believe the most plausible interpretation of our data is that the predictive codes fed back to V1 from V5 update to new retinotopic post-saccadic locations and amplify visual responses when they do not fit the temporal prediction triggered by the surrounding apparent motion illusion(Alink et al., 2010; Vetter et al., 2015). As V1 is acallosal (Dumoulin and Wandell, 2008; Saenz and Fine, 2010; Van Essen et al., 1982) predictive feedback must relocate via higher cortical areas. hMT/V5+, as a motion sensitive region (Chawla et al., 1998; Goebel et al., 1998) known to integrate long-range apparent motion (Muckli et al., 2005, 2002; Sterzer et al., 2006b; Vetter et al., 2015; Wibral et al., 2009), is a likely candidate. From our fMRI data, we do not know the temporal dynamics of the prediction relocation (see Vetter et al., 2012), but importantly, we do know it is effective in time for the post-saccadic processing.

## Conclusion

Predictive coding theories of vision state that cortical predictions are compared with sensory signals to compute an error in the case of mismatch. These error signals update the brain’s internal models of the visual world to stabilize perception, cognition and behavior. However, with each saccade, the visual system is flooded with a new pattern of sensory signals. Therefore, to be functionally useful, predictions must feed back to the relevant retinotopic location at which their sensory counterpart will arrive to V1 after each saccade. We show that predictions do update retinotopic position in time for post-saccadic input to V1. Hence, predictive coding is an ecologically viable mechanism of the human visual system.

## Methods

### Subjects

We recruited 27 healthy subjects with normal or corrected-to-normal vision (15 male; 19-34 years) using the University of Glasgow, School of Psychology subject pool. Subjects gave written informed consent. The ethics committee of the College of Science and Engineering, University of Glasgow provided ethical approval. We performed a power analysis based on data from Alink et al., (2010) where subjects performed a very similar apparent motion task in the fMRI, but without eye-movements. The power analysis estimated a minimum sample size of n=5 (effect size = 0.06, standard deviation = 0.04, null hypothesis = 0, type I error rate = 2.5%, power = 0.8). We then performed a pilot experiment in which we combined the apparent motion effect with eye-movements. The subsequent power-analysis of five pilot-subjects confirmed the effect size from Alink et al., (2010) (see supplemental material, effect size = 0.08, standard deviation = 0.05, null hypothesis = 0, type I error rate = 2.5%, power = 0.8). For the main experiment we settled for a group size of 10, well above the estimated minimum group size for these effects. Due to the saccade criterion (see further in methods), one subject was removed in the Main apparent motion experiment (leaving n = 9), two subjects were removed from the flicker control experimental analysis (leaving n = 5), and two subjects were removed from the analysis of the Replication experiments (leaving n = 8). Two subjects who took part in the flicker control also performed the replication experiments (CV & GV).

### Apparent Motion Stimulus

We presented the apparent motion stimulation in the center of a grey screen (RBG: 153,153,153; Figure 1). We induced the illusion of vertical motion by flashing two white rectangles in an alternating rhythm at a frequency of 3.75 Hz (Figure 1). These two white rectangles were either 14.84° (pilot experiment) or 8.84° of visual angle apart from each other (main experiment). Each rectangle was presented for 5 frames (83.3ms), followed by an inter-stimulus interval of 5 frames. In the pilot experiment, we presented a green fixation cross (2.1° visual angle in size) 10.27° to the left of the apparent motion illusion centre, and a red cross (sized 2.1°) 10.27° to the right of the illusion. In the main fMRI experiment, the fixation crosses were 0.7° size and positioned 5.81° horizontally from the illusion. Subjects fixated on the red cross. The red cross alternated horizontally with the green cross prior to the 7^th^ cycle of apparent motion, approximately 2 s after the apparent motion onset, cuing subjects to saccade across the illusion (Figure 1a-d). Shortly after the subjects’ saccade landed, a target appeared (for one frame of 16.67ms) during the 8^th^ cycle and the illusion ceased. This design ensured that the right hemisphere processed the illusion (and a prediction was generated in higher visual areas) while the left hemisphere processed the target. Targets presented on the apparent motion trace were either in-time or out-of-time with the illusion. In-time targets were presented approximately 83 ms after saccade offset and out-of-time targets approximately 50 ms after saccade offset (which was 117 ms and 150 ms into the 8^th^ cycle respectively). There were three apparent motion conditions: in-time target, out-of-time target and no target. Our previous data revealed that subjects take approximately 300ms between saccade cue and saccade completion(Vetter et al., 2012), hence we presented the target approximately 450ms after the red fixation cross shifted location, in the 8^th^ cycle of apparent motion. After the 8^th^ cycle, subjects were presented with the question ‘target?’ indicating that they should respond ‘yes’ or ‘no’ via button-press. Subjects had to respond within 1500ms, there was then an inter-trial-interval of 1333ms prior to the beginning of the next trial. In total 6 conditions were presented: 60% apparent motion conditions (3 conditions of 20% of each), 20 % of the mapping conditions (2 conditions, 10 % each), and 20% of the baseline condition which was the presentation of the fixation and alternating cross alone, without any saccade required. Our replication apparent motion experimental design had no alterations from the original design.

### Mapping Stimulus

We presented mapping conditions in the same runs as the apparent motion illusion. Mapping conditions comprised static images of the lower apparent motion inducer and the target, and were both presented for 4000 ms (Figure 1b). These conditions enabled us to map the separate locations in V1 that respond to the inducing stimuli and the target (GLM contrast of Target >Lower inducing stimulus: beta-weight = 0.88, t= 18.53, p<0.0001 in the main fMRI experiment; beta-weight (SE)=0.92(0.06), t=18.81, p<0.0001 in the flicker control experiment; beta-weight (SE)=1.139(0.043), t=26.221, p<0.0001 in apparent motion replication experiment; beta-weight (SE)=4.313(0.185), t=23.368, p<0.0001 in flicker control replication experiment).

### Flicker control stimulus

We tested five new subjects in an fMRI experiment. The flicker control stimulation was identical to the apparent motion stimulation apart from the two blinking inducer stimuli were presented simultaneously instead of in alternating rhythm. The upper and lower inducers were presented in unison for 5 frames (83.3ms), followed by an inter-stimulus interval of 5 frames, resulting in a flicker at the same rate as the apparent motion stimulus (3.75 Hz; Figure 1e). All other parameters remain the same as the main fMRI experiment. We cued subjects to saccade across the flicker stimulus and we presented targets after the saccade in the final flicker cycle, with the same timings as in the main experiment. The flicker stimulus was processed by the right V1 and the targets were presented to the left V1. No design alterations were made during our replication of the flicker control experiment. *Procedure.* Subjects completed four functional runs of 15 minutes each. In the main fMRI, flicker control, and replication experiments, subjects viewed stimuli on an fMRI compatible screen positioned in the bore of the magnet at a distance of 110 cm (screen resolution: 1024 x 768). We used Neurobehavioral systems Presentation^®^ software (Version 14.9) to present the paradigm, with a screen refresh rate of 60Hz.We presented the three apparent motion conditions and a baseline condition of 25 times per run and 100 times across the whole experiment. We presented the two mapping conditions 12 times each per run and 48 times across the experiment. We used a randomization scheme to order the trials, ensuring that no triplets of conditions were repeated.

### MRI Data Acquisition

We acquired functional and anatomical MRI data using a 3 Tesla MRI system (Siemens Tim Trio) with a 12-channel head coil (and a 32-channel head coil for the control and replication experiments). For the functional scans, an echo-planar imaging sequence had the following parameters: 17 slices, TR-1, TE-30, 860 volumes per run, an FOV of 205 mm, and a resolution of 2.5 x 2.5 x 2.5 mm. The 17 slices were orientated perpendicular to the calcarine sulcus to capture the early visual cortex. The anatomical MRI sequence used had a TR of 1.9, 192 volumes, and a resolution of 1 x 1 x 1 mm. fMRI parameters and analyses were identical in main, pilot, flicker control, and replication experiments.

### Eye-tracking Acquisition

In the main experiment and flicker control, subjects’ eye-movements were recorded using an Eyelink 1000 (SR Research) mounted on the fMRI compatible projector screen with a sampling rate of 500Hz (calibrated at the start of each run). The data was recorded by Eyelink software and downloaded for analysis using Eyelink Data Viewer.

### Saccade Criterion

The saccade criterion denoted that subjects had to complete a saccade 400 ms after cue, and the saccade must cover at least 200 pixels horizontally across the apparent motion between onset and offset (Figure S5a). The criteria ensured that subjects processed the apparent motion in the right hemisphere and the target in the left. Specifically, in-time and out-of-time targets were presented at least 83ms and 50ms after latest saccade offset respectively, which was 483 and 450ms after cue to saccade respectively. Subjects had to land their saccade between the cross color change + 333ms and 400ms (i.e. within a 67ms time-window), to ensure that trials are outside the saccadic suppression window of +/-150ms. We excluded trials where a saccade did not meet the criterion. In the main fMRI experiment one subject and one run from two other subjects who showed less than 20 trials per run with a correct saccade were excluded (Figure S5b). Two subjects were excluded from the flicker control experiment as the saccade criterion reduced their successful saccade trials to less than 20 per run. We removed a further two subjects and 1 run from one other subject due to less than 20 successful trials per run in the replication experiments.

### MRI Analysis

The functional and anatomical data were analyzed using Brainvoyager QX® software (Version 2.4). We discarded the first two volumes of each functional run to avoid saturation effects. To remove low-frequency noise and drift, we performed high-pass filtering at 6 sines/cosines during the 3D-motion correction for each run. After preprocessing, we aligned the functional data with the high-resolution anatomical data and transformed into Talairach space (applying each subject’s brain in to a common space along the AC-PC plane). We created 3D aligned time courses (VTCs) for each run after the intra-session anatomical was aligned with the high resolution anatomical. Each subject’s cortex was inflated into a surface model using the manually inhomogeneity corrected high resolution anatomical.

Once we removed invalid eye-tracking trials, we performed single subject and group deconvolution analysis. We contrasted beta-weight values over 3-7 seconds after stimulus onset in the target processing region and apparent motion inducer processing region of left V1 separately, corresponding to the time when the targets were presented in the apparent motion trials compensating for BOLD lag. We performed the same analysis in the left V2, right V5, and left V5. We normalised our data to baseline (fixation only) before performing our analyses. We contrasted the response amplitudes (expressed in beta-weight values of a general linear model (GLM) analysis) for in-time and out-of-time target trials using a serial correlation corrected comparison to determine the activation difference. Response amplitudes were also contrasted for flicker with in-time targets and flicker with out-of-time targets in the control analysis. We used a deconvolution analysis in order to analyze the BOLD responses between different conditions. The same analyses were performed in the replication experiments.

### Retinotopically defining regions of interest (ROI)

The region of interest in left V1 was identified using a GLM contrast of target>lower inducing stimulus (mean (SD) Talairach co-ordinates: *x* = −11.33 (4.4), *y* = −89.67 (2.7), *z* = −6.3 (7.5); FDR = 0.05; Figure 2). Left V2 was defined using the target>lower GLM contrast (mean (SD) Talairach co-ordinates: *x* = −18.11 (5.6), *y* = −94 (3.9), *z* = 1.89 (6.4); FDR = 0.05). The ROIs for right V5 and left V5 and right V1 were defined using apparent motion without target condition > baseline GLM contrast as no mapping data were collected for these regions (mean (SD) Talairach co-ordinates for right V5: *x* = 43.22 (4.4), *y* = −65.78 (4.1), *z* = 0.89(5.3); left V5: *x* = −45.22 (6.0), *y* = −70.44 (3.2), *z* = −0.89 (4.2); right V1: *x* = 7.78 (3.3), *y* = −80.89 (6.9), *z* = −1.78 (7.0); FDR = 0.05). The apparent motion inducer processing region of left V1 was also selected for control analysis using lower inducer > target contrast (mean(SD) Talairach co-ordinates: *x*= −5.89(5.3), *y* = −90.67 (6.2), *z*= 0.22(6.9) ; FDR = 0.05). For the flicker experiment, the target processing region in V1 was also defined using target>lower inducing stimulus (mean (SD) Talariach co-ordinates: *x* = −11.6 (4), *y* = −86.4 (3.4), *z* = −5.2 (7.2). We defined all ROIs with the same contrasts in the replication experiments.

### Behavioral Analysis

We recorded behavioral data using Neurobehavioral System’s Presentation® software during the fMRI runs. We excluded one subject in the main fMRI experiment due to data recording issues. We conducted the analysis after applying the saccade criteria. Target detection accuracy was averaged per subject and then across all subjects. Analysis focused on accurate detection difference of in-time versus out-of-time targets.

### Psychophysical control experiment

It was our objective to have the target prediction effect generated by pre-saccadic processing and not by post saccadic, post-diction mechanisms. For this reason, the target always appeared in the post saccadic stimulated right visual hemi-field, directly after saccade. The pre-saccadic illusion perception primed the visual system for prediction effects prior to subjects performing the saccade. After subjects executed the saccade, the post-saccadic stimulated hemisphere processed only the target and the lower apparent motion inducer. It is the top-down cross-hemispheric prediction of motion that leads to the difference between in-time and out-of-time target presentations. In 20% of trials, no target appeared but in 40% of the trials an in-time or an out-of-time target (20%/20%) appeared, allowing us to measure brain-activity differences between these conditions. In the remaining 40% of trials, we presented the mapping conditions and baseline. The target trials (40%) were effective in inducing different BOLD activations but did not create any behavioral effects in terms of target detection. This may be because subjects knew the target would be presented after saccade. We have shown robust behavioral effects previously (Alink et al., 2010; Schwiedrzik et al., 2007; Vetter et al., 2015, 2014, 2012) using a paradigm in which targets were also presented mid-fixation (thus not always after saccade). We aimed to replicate this behavioral detection advantage in the current experiment. We ran an extra session behavioral experiment using a continuous apparent motion paradigm to enable target presentation both after saccade and during fixation.

### Subjects

Nine subjects (6 male; 19-28 years) who participated in fMRI experiment 2 also completed the psychophysical counterpart. We excluded one of these subjects from analysis using the same saccade criterion used for trial exclusion during fMRI (Figure S5). The ethics committee of the College of Science and Engineering, University of Glasgow provided ethical approval.

### Stimuli

We presented the paradigm using Neurobehavioral Systems’ Presentation® (Version 14.9) software, using identical parameters for the three apparent motion conditions as in the fMRI experiments (see *Apparent Motion Stimulus*). However, the apparent motion stimulation was continuous (onset and offset of each trial were not detectable) and no mapping conditions were presented. Subjects were cued to saccade across the apparent motion illusion every 2.66 s trials consisted of 10 cycles of apparent motion, and targets were presented in the cycle directly after saccade (at the same time as in the fMRI experiment) or during fixation.

### Protocol

We presented the three apparent motion conditions at random within 5 runs of the experiment. In 60% of the 152 trials per run, we presented targets directly after saccade in the same time-period as was used for the fMRI experiment. We presented a target mid-trial during fixation in 20% of trials, and in the remaining 20% of trials we did not present a target. This design decreased the probability of targets appearing after the saccades. Every 25 trials, the apparent motion was interrupted for 20 s with a natural scene to allow subjects to rest their eyes and prevent apparent motion breakdown (Anstis et al., 1985).

### Task & Procedure

Subjects were seated 70 cm from a 16 inch Sony Trinitron CRT Monitor (1024 x 768; 60Hz), upon which we presented the stimuli. Subjects’ heads were supported using chin and forehead rests. We recorded subjects’ eye-movements continuously throughout the experiment (EyeLink 1000, SR Research; acquisition as in fMRI method). We instructed subjects to fixate on the red fixation cross and move their eyes across the illusion when the red cross alternated with the green. We instructed subjects to indicate target detection with a button press.

### Analysis

The behavioral data was recorded using Presentation® software. We applied the same eye-tracking criterion for the fMRI data analysis to each trial of the psychophysical experiment. Detection of targets was only included if the button press occurred within 150 and 1200 ms after target onset. Target detection accuracy was averaged per subject and then across all subjects. Analysis focused on accurate detection difference of in-time versus out-of-time targets.

## Acknowledgements

This work was supported by BBSRC grant (BB/G005044/1), ERC grant StG 2012_311751-Brain reading of contextual feedback and predictions’) and Human Brain Project grant (‘Context-sensitive multisensory object recognition: a deep network model constrained by multi-level, multi-species data’).

